# Humans can learn bimodal priors in complex sensorimotor behaviour

**DOI:** 10.1101/2025.02.12.637788

**Authors:** Stephan Zahno, Damian Beck, Ernst-Joachim Hossner, Konrad Kording

**Affiliations:** Institute of Sport Science, University of Bern, Switzerland; Department of Neuroscience, University of Pennsylvania, USA

**Keywords:** Bayesian integration, probabilistic inference, complex tasks, naturalistic behaviour, sensorimotor control

## Abstract

Extensive research suggests that humans integrate sensory information and prior expectations in a Bayesian manner to reduce uncertainty in perception and action. However, while Bayesian integration provides a powerful explanatory framework, the question remains as to what extent it explains human behaviour in naturalistic situations, including more complex movements and distributions. Here, we examine whether humans can learn bimodal priors in a complex sensorimotor task: returning tennis serves. Participants returned serves in an immersive virtual reality setup with realistic movements and spatiotemporal task demands matching those in real tennis. The location of the opponent’s serves followed a bimodal distribution. We manipulated visual uncertainty through three levels of ball speeds: slow, moderate, and fast. After extensive exposure to the opponent’s serves, participants’ movements were biased by the bimodal prior distribution. As predicted by Bayesian theory, the magnitude of the bias depends on visual uncertainty. Additionally, our data indicate that participants’ movements in this complex task were not only biased by prior expectations but also by biomechanical constraints and associated motor costs. Intriguingly, an explicit knowledge test after the experiment revealed that, despite incorporating prior knowledge of the opponent’s serve distribution into their behaviour, participants were not explicitly aware of the pattern. Our results show that humans can implicitly learn and utilise bimodal priors in complex sensorimotor behaviour.

## Introduction

Due to noise and delays in the nervous system, humans perceive the world and act upon it under considerable uncertainty^1,2^. Dealing with uncertainty is a fundamental challenge for human behaviour^3,4^. A functional strategy to reduce uncertainty, formalised by Bayesian integration, is to combine current sensory information with prior expectations and weighting both sources according to their reliability^5,6^. Extensive research demonstrates that human behaviour is consistent with Bayesian integration across lab tasks such as reaching^7^, pointing^8^, as well as estimating speeds^9^, forces^10^ and timings^11^. Over the last two decades, Bayesian theory has become highly influential as a unifying framework for perception^12^, higher-level cognition^13^, and motor control^5^. However, despite its explanatory power in controlled experimental settings, an important question remains: To what extent does Bayesian integration explain behaviour in the complex situations encountered in our daily lives^14^?

In recent years, testing major theories of human behaviour in complex, naturalistic tasks has been increasingly highlighted as a key challenge in psychology and neuroscience^15–20^. On closer examination, the demands in classical lab tasks differ from many real-world tasks in two regards. First, in most experiments, striving for high experimental control, subjects remain seated and are asked to react to abstract stimuli with simple arm movements or button presses in a highly constrained manner. Many real-world tasks—think of returning a tennis serve— require coordinating complex movements. Only recently, an increasing number of studies have begun to take on the challenge of testing Bayesian predictions in more complex movement tasks^21–25^. Second, in most experiments on Bayesian integration, participants learn regularities in the task that follow a simple Gaussian distribution, whereas many real-world tasks contain more complex patterns of regularities. Again, tennis provides a natural example: Typically, the serves of your opponent are not normally distributed around one location but rather contain two peaks of highly probable ball locations—a wide vs. a T serve^26^—following a bimodal distribution. However, while Bayesian models make clear predictions on how bimodal priors should be integrated^7^ and, thus, provide a hard test case for Bayesian predictions, the extent to which humans can learn and exploit bimodal priors has been addressed only in a few studies and remains a subject of debate^7,27,28^.

Here, we investigate whether humans can learn and use bimodal priors in a complex sensorimotor task—that is, the task of returning a tennis serve. We developed an immersive virtual reality (VR) setup that enables the study of natural movement behaviour with spatial and temporal constraints that match real tennis while simultaneously ensuring full experimental control. In three experimental sessions, participants had the task of returning tennis serves. The locations of the opponent’s serve followed a bimodal distribution, with peaks at 50 cm and 90 cm to the right of the participants’ starting position (Figure 1a). We did not inform the participants about the pattern of the opponent’s serves. We manipulated the uncertainty of visual information by varying among three levels of ball speeds: slow, moderate and fast (i.e. ball trajectories that are easy vs. difficult vs. very difficult to perceive). As a measure of the participants’ estimation error, we computed the horizontal deviation between the ball and the racket’s sweet spot (i.e. the centre of the racket) at the time of contact.

**Fig. 1.**
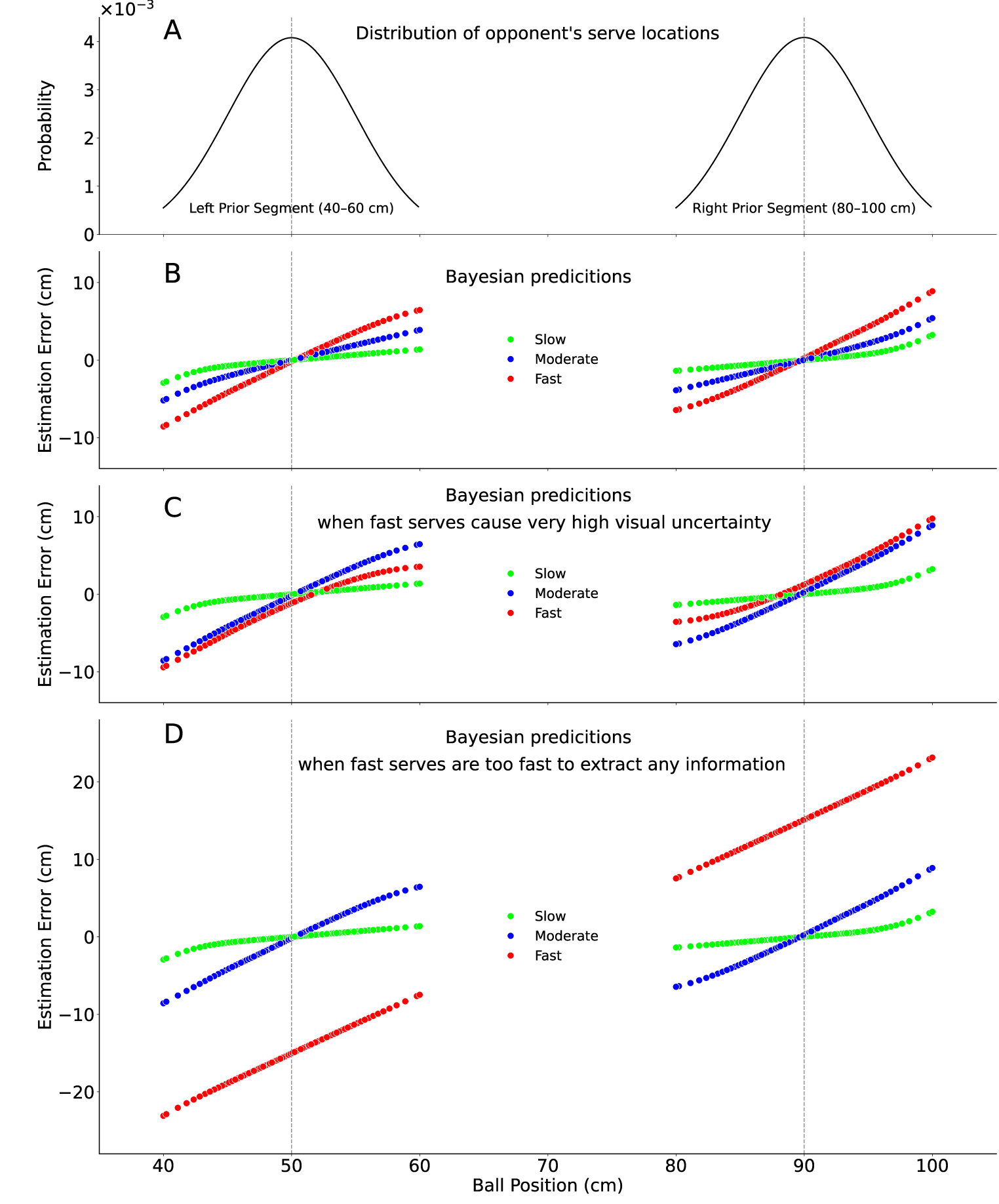
Expectations based on Bayesian simulation. **A**: The plot shows the distribution of the opponent’s serve locations. The ball positions range from 40–60 cm and 80–100 cm to the right of the participant’s initial position. The peaks of the distributions are at 50 cm and 90 cm (SD = 5 cm). **B:** We ran a Bayesian simulation^29^ with three different levels of uncertainty (slow, moderate, and fast ball speeds) and a bimodal prior distribution. The plot depicts the simulated estimation errors (difference ball – racket’s sweet spot) (y-axis) in relation to the true ball positions (x-axis). In this example, the standard deviation of the likelihood is set to 2 cm for slow, 4 cm for moderate, and 10 cm for fast serves. The simulation code is available on GitHub (https://github.com/ispw-unibe-ch/bt-bimodal_prior_integration_vr_tennis). **C:** This plot depicts a simulation in which we assume that the ball speed has a higher effect on visual uncertainty. The standard deviation of the likelihood is set to 2 cm for slow, 10 cm for moderate, and 14 cm for fast serves. The simulation shows that when uncertainty increases to very high levels, the relationship between error and ball positions becomes more linear. **D:** This plot illustrates the extreme example of when fast serves are so fast that visual inputs become uninformative. The standard deviation of the likelihood is set to 2 cm for slow, 10 cm for moderate, and 40 cm for fast serves. In this extreme case, the model predicts that the error linearly depends on the true ball position with a slope of 1.

To test whether participants learned the bimodal prior and integrated it with sensory information according to Bayesian principles, we analysed the relationship between the estimation error and the true ball location analogous to the analyses in classical Bayesian experiments^7^. Figure 1 (b-d) depicts Bayesian predictions with a bimodal prior for the three uncertainty conditions: slow (green), moderate (blue), and fast (red) serves. Higher ball speeds are assumed to cause higher visual uncertainty. Generally, the Bayesian model predicts that participants’ movements are biased towards the prior and that the magnitude of the bias depends on visual uncertainty. Specifically, for the bimodal case, the model predicts a non-linear relationship between the estimation error and the true ball location, which can be quantified with a ‘jump’ between the left and right segments of the distribution. Throughout the paper, we refer to this jump as the ‘bimodal prior effect’. Based on Bayesian integration, we expect that, after extensive exposure to the opponent’s serve distribution, (1) a bimodal prior effect can be detected and (2) the magnitude of the bimodal prior effect depends on visual uncertainty. Bayesian simulations with different levels of uncertainty illustrate that the bimodal prior effect should increase with higher uncertainty (Figure 1b); however, this happens only up to a certain threshold. When fast serves cause very high uncertainty, the relationship becomes more linear again (Figure 1c). In a (hypothetical) extreme case where balls are so fast that no information can be extracted, the model predicts that participants’ responses shift towards the mean of the distribution independent of the true ball position, thereby resulting in a linear relationship between estimation error and true ball location (Figure 1d). The results of our experiment provide evidence that humans can learn a bimodal prior in complex sensorimotor behaviour and utilise it in a Bayesian manner.

## Results

The task of the 24 right-handed participants was to return tennis serves towards the centre of a target on the opponent’s field in a customised life-sized VR CAVE environment (see video on GitHub https://github.com/ispw-unibe-ch/bt-bimodal_prior_integration_vr_tennis). On three days, participants returned 1,440 serves that followed a bimodal distribution (mixture of Gaussians). The serve locations were distributed in a range between 40–60 cm and 80–100 cm to the right of the participants’ starting position (Figure 2, right). Importantly, while the trajectory of the approaching ball varied from trial to trial, the body movements of the serving avatar were identical for all serves. In other words, the opponent’s serving motion was uninformative and provided no cues regarding the ball’s trajectory. This design ensured that the participants could rely only on two sources of information: current sensory input regarding the ball trajectory and prior knowledge accumulated from previous trials. We manipulated visual uncertainty by varying the initial ball speeds across three levels: slow (v = 108 km/h), moderate (v = 180 km/h), and fast (v = 252 km/h). The serves followed a prespecified spline trajectory (Figure 2, left), and we controlled for the temporal timing demands to be the same in all speed conditions. Unbeknownst to the participants, they were actually striking an invisible stationary ball at a fixed location along the trajectory. This stationary ball could be hit within a 400 ms window—maximally 200 ms before and 200 ms after the visibly displayed ball passed its location. The deviation between the racket’s sweet spot at impact and the position of the invisible ball determined the direction of the return and, thus, directly affected task performance (Figure 2, right). After each return, points were displayed on a scale of 0– 100 to indicate how well the participant hit the ball.

**Fig. 2.**
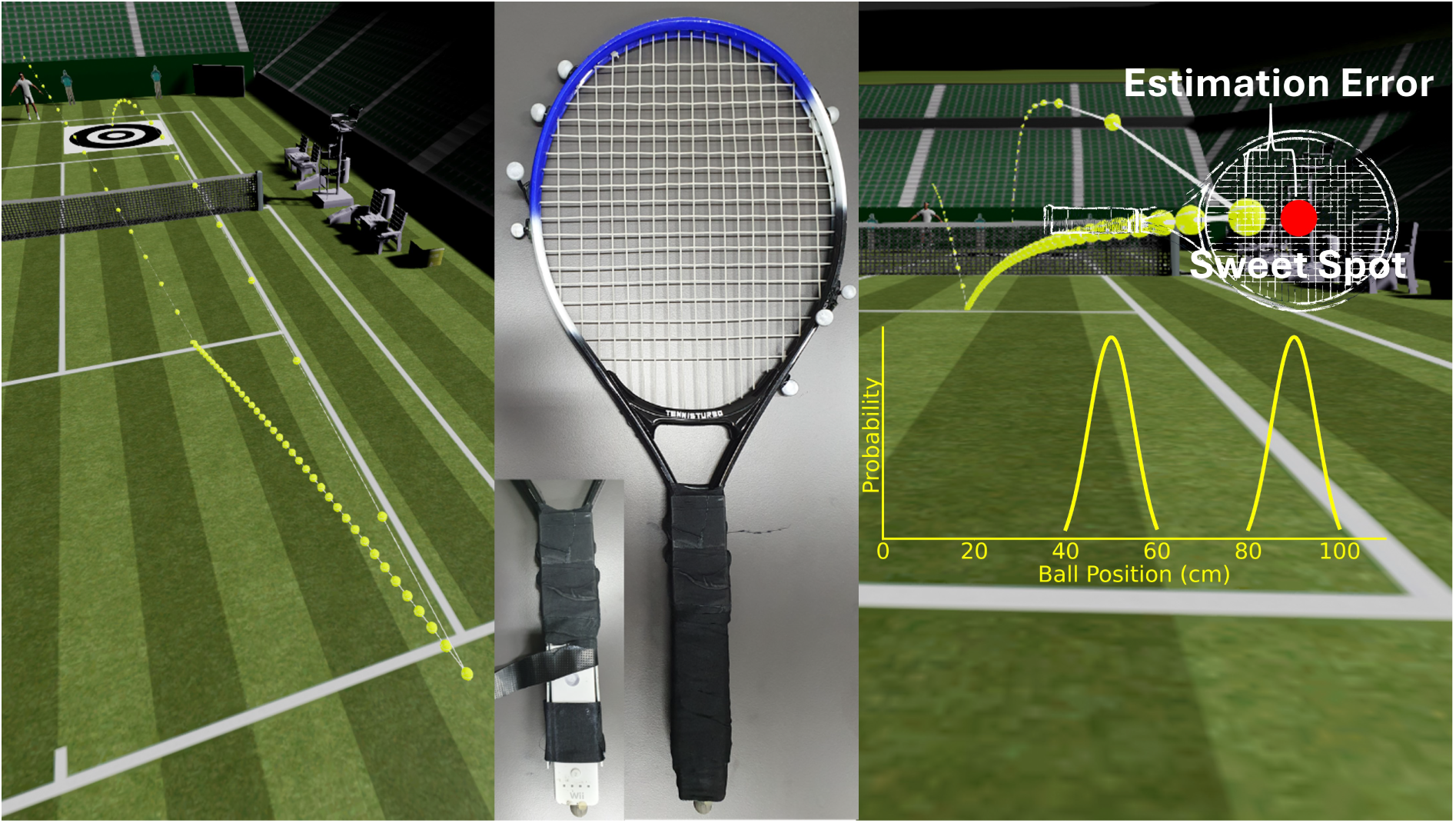
Experimental Virtual Reality Setup. **Left:** The approaching ball followed a prespecified spline trajectory, and the participants’ task was to return the ball towards the centre of the target on the opponent’s court. **Centre:** Participants swung a real tennis racket with an integrated Wii for haptic feedback vibration and placed markers to track the racket’s location. **Right:** The direction of the return trajectory was determined by the estimation error. A positive estimation error implies that the sweet spot is on the left of the ball (i.e. the participant does not reach far enough to the right). A negative estimation error implies that the sweet spot is on the right of the ball (i.e. the participant reaches too far to the right). In this example, the estimation error is negative. If the ball was hit perfectly with the sweet spot (zero estimation error), the return was perfectly in the direction towards the centre of the target, and the maximum score of 100 was displayed.

To test whether participants can learn a bimodal prior and utilise it in a Bayesian manner, we examined the relationship between the estimation error and true ball positions. According to the Bayesian model, participants’ estimation error should be biased by the bimodal prior as a function of uncertainty. This bias results in a non-linear relationship between estimation error and true ball positions, which we quantify by a ‘jump’ between the left and right sides of the distribution (bimodal prior effect). To examine the bimodal prior effect, we calculated multilevel regression analyses with two independent variables: true ball position and a bimodal segment factor (left vs. right). The same analysis was conducted for the three uncertainty conditions. Based on the assumption that the acquired experience regarding the bimodal prior needs 24 hours to be sufficiently consolidated during one night of sleep^30^, we separately analysed all returns for each condition on day 1 (initial practice phase; Figure 3, right) and all returns for each condition on days 2 and 3 together (after extensive practice; Figure 3, left).

**Fig. 3.**
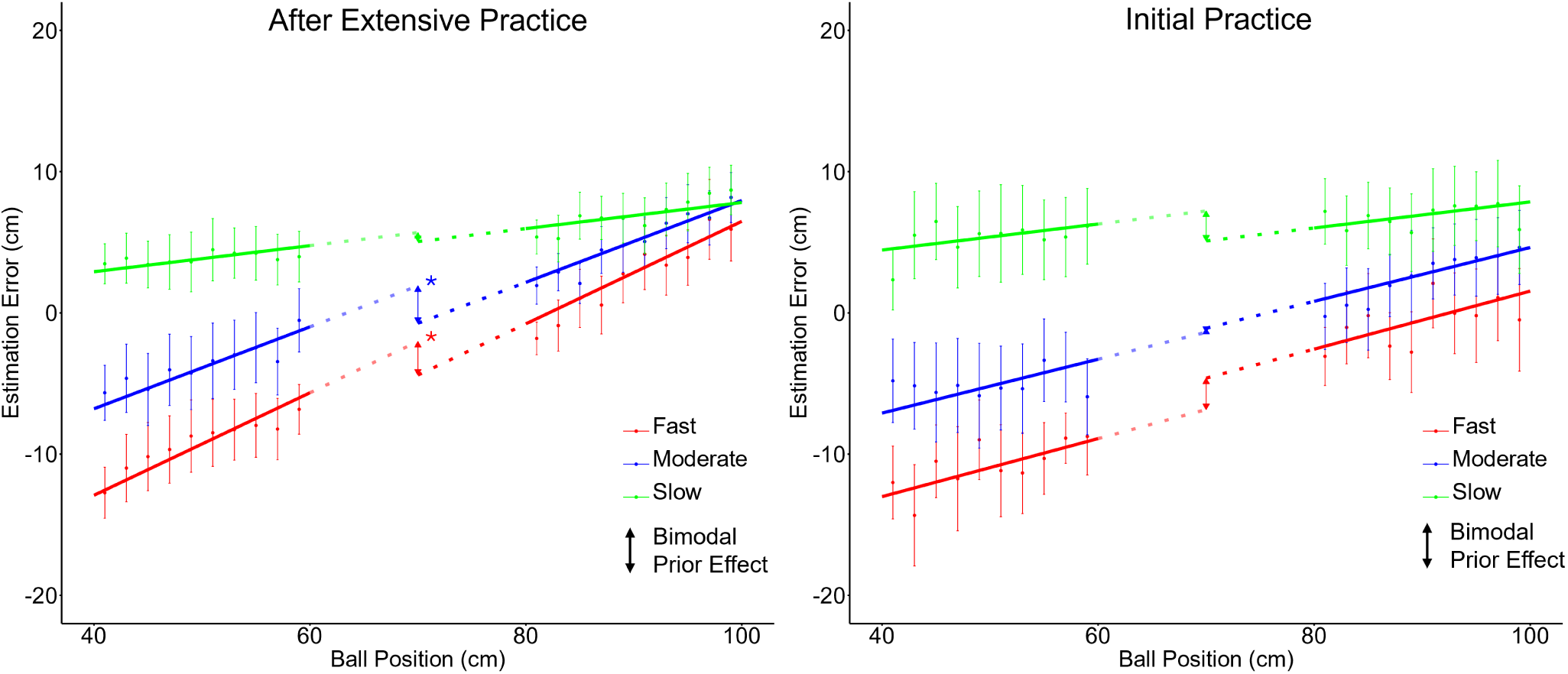
Bimodal prior effect as a function of visual uncertainty. **Left:** The graph displays the estimation error (difference ball – racket’s sweet spot) as a function of the true ball position and the uncertainty condition (slow, moderate, fast) after extensive exposure to the opponent’s serves (data from days 2 and 3). The data are divided into 10 bins on each side. For each bin, the arithmetic mean and 95% confidence interval are displayed. The multilevel regression model is computed with the independent factors of ball position and bimodal segment factor (left 40–60 cm, right 80–100 cm). The ‘jumps’ between the left and right sides of the distribution quantify the bimodal prior effect. **Right:** The same plot is depicted for the initial acquisition phase (data from day 1).

### Participants learn and use the bimodal prior

Consistent with Bayesian predictions, after extensive exposure to the opponent’s serves, non-linear effects were found (Figure 3, left). The magnitude of the bimodal prior effect depends on the uncertainty condition. Significant bimodal prior effects were found for fast serves (red) (*N*_subjects_ = 24, *N*_measurements_ = 5597*, B* (0 = left, 1 = right) = –2.36 [–4.39, –0.33], *t*(5571) = – 2.28, one-sided and Bonferroni-Holm corrected *p* = .023) and for moderate serves (blue) (*N*_subjects_ = 24, *N*_measurements_ = 6610*, B* (0 = left, 1 = right) = –2.64 [–4.43, –0.86], *t*(6584) = – 2.90, one-sided and Bonferroni-Holm corrected *p* = .006). No bimodal prior effect was observed in the slow condition (green) (*N*_subjects_ = 24, *N*_measurements_ = 6488*, B* (0 = left, 1 = right)= –0.81 [–2.38, 0.76], *t*(6462) = –1.02, one-sided and Bonferroni-Holm corrected *p* = .154) (for full statistics and model comparisons, see Tables 1–6 in Extended Data). This pattern of results aligns well with the core predictions derived from the Bayesian model. It suggests that subjects acquired the bimodal prior and relied on the learned distribution as a function of uncertainty.

### Learning a bimodal prior requires extensive practice

On day 1, no evidence for a bimodal prior effect was found (Figure 3, right). Across all speed conditions, the relationship between error and true ball position was linear. The regression analyses indicate no significant bimodal prior effects for the fast (red) (*N*_subjects_ = 24, *N*_measurements_

= 2699*, b* (0 = left, 1 = right) = 3.24 [–0.01, 6.49], *t*(2673) = 1.95, two-sided and Bonferroni-Holm corrected *p* = .153), moderate (blue) (*N*_subjects_ = 24, *N*_measurements_ = 3122*, b* (0 = left, 1 = right) = 0.27 [–2.60, 3.15], *t*(3096) = 0.19, two-sided and Bonferroni-Holm corrected *p* = .852), nor for the slow condition (green) (*N*_subjects_ = 24, *N*_measurements_ = 3075*, b* (0 = left, 1 = right) = – 2.24 [–4.97, 0.49], *t*(3049) = –1.61, two-sided and Bonferroni-Holm corrected *p* = .216) (for full statistics and model comparisons, see Tables 7–12 in Extended Data). These results indicate that participants had not yet learned the bimodal prior by day 1. This finding aligns with previous research showing that acquiring complex distributions requires extensive practice^7,30^.

### Biomechanics additionally bias participants’ movements

In addition to the bimodal prior effect, data show a general positive linear trend between the estimation error and ball position (Figure 3, left and right). This effect can be explained by biomechanical considerations. A ball position of approximately 90 cm from the body enables participants to swing the racket with an almost stretched arm, thereby resulting in low motor costs. In contrast, when the ball is played closer to the body, participants must flex their arm or take a small step to the left, which incurs higher motor costs. Based on established findings in the motor control literature^31–34^, we expected that participants’ movements would be generally biased towards positions with lower motor costs. This would result in a predictable pattern of errors: When the ball is played close to the participants’ body, they are more likely to hit too far to the right, leading to negative errors, particularly when the balls are fast.

To test this biomechanical explanation, we conducted a control experiment with 24 new participants. The task was identical to the main experiment, but serve locations followed a uniform distribution (equally spread across the complete range of 40–100 cm) rather than a bimodal distribution. This enabled us to isolate errors caused by biomechanical constraints rather than prior expectations. Figure 4 (left) depicts the results of the control experiment. As expected when taking biomechanical costs into account, the relationships between estimation error and true ball positions are linear, with steeper slopes for higher speed conditions and an intersection at approximately 90 cm. The regression analyses indicate significant slope effects for the fast (red) (*N*_subjects_ = 24, *N*_measurements_ = 2763*, B* (ball position) = 0.38 [0.31, 0.45], *t*(2738) = 10.80, two-sided and Bonferroni–Holm corrected *p* < .001) and moderate conditions (blue) (*N*_subjects_ = 24, *N*_measurements_ = 3043*, B* (ball position) = 0.15 [0.09, 0.21], *t*(3018) = 4.93, two-sided and Bonferroni-Holm corrected *p* < .001), but not for the slow condition (green) (*N*_subjects_ = 24, *N*_measurements_ = 2919*, B* (ball position) = –0.05 [–0.09, 0.00], *t*(2894) = –1.96, two-sided and Bonferroni-Holm corrected *p* = .050) (for full statistics and model comparisons, see Tables 13–18 in Extended Data). Figure 4 (right) illustrates the motor bias in participants’ average movement trajectories. When the opponent served at locations of approximately 90 cm (bin: 87–93 cm), the trajectories were unbiased, while trajectories at 50 cm (bin: 47–53 cm) showed a rightward bias. This bias increased with higher ball speeds. These data confirm that the positive linear trend in Figure 3 (left and right) can be attributed to biomechanical constraints, while the non-linear effects (i.e. the ‘jumps’) can only be explained by the bimodal prior.

**Fig. 4.**
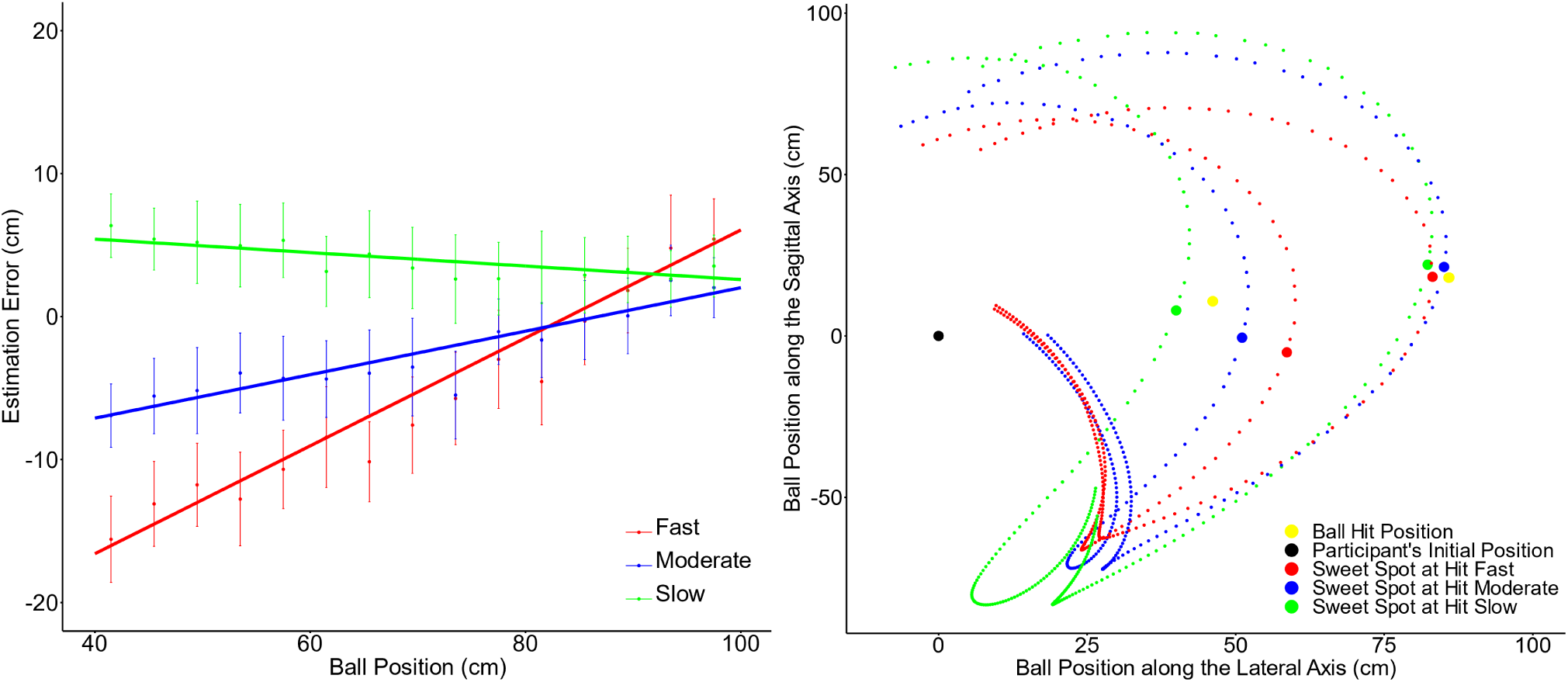
Biomechanical effect in the control experiment. **Right:** The graph displays the estimation error (difference ball – racket’s sweet spot) as a function of the true ball position and the uncertainty condition (slow, moderate, and fast) in the control experiment. The graph depicts systematic errors that are produced by biomechanical factors. The data are divided into 15 bins. For each bin, the arithmetic mean and 95% confidence interval are displayed. **Left:** The plot illustrates the trajectories of the racket’s sweet spot (averaged across participants) for serve locations at approximately 50 cm (bin: 47–53 cm) and approximately 90 cm (bin: 87–93 cm) from a top-down perspective.

### Prior knowledge of the bimodal distribution is implicit

After the final experimental session, we asked participants if they recognised a pattern in the opponent’s serve distribution. On a schematic tennis court on paper, no participant was able to draw hit positions split into left and right parts or anything similar to a bimodal distribution. In the following forced-choice question (see Appendix), we asked participants to choose between four potential distributions: uniform, normal (unimodal), bimodal, or a scenario in which the ball always landed in the same location. Only 6 out of 24 participants crossed the bimodal distribution; just as one would expect to accidentally occur by chance with four answer choices. Fifteen participants crossed the normal distribution, with the highest probability in the centre where the ball actually never landed. This result suggests that participants integrated implicit prior ‘knowledge’ of the bimodal distribution into their tennis return, without any explicit awareness of the pattern.

## Discussion

In this study, we investigated whether humans can learn bimodal prior distributions and use them according to Bayesian principles in complex sensorimotor behaviour. Over three days, participants returned 1,440 tennis serves that followed a bimodal distribution. Our results reveal that, through playing tennis, participants acquired a bimodal prior and used it in action. Specifically, after extensive experience, we found a non-linear relationship between the estimation error and the true ball position—a ‘bimodal prior effect’—that depends on uncertainty, as predicted by Bayesian theory^7^. Remarkably, despite being exposed to the opponent’s serving pattern over 1,440 trials and incorporating this information into their behaviour, participants were unable to report, above chance level, whether the opponent’s serves followed a bimodal distribution when asked explicitly. This finding highlights that the sensorimotor system can utilise prior knowledge of environmental statistics to optimise behaviour without the need for explicit awareness of these patterns.

While our results align well with the core predictions derived from the Bayesian model^29^, the data clearly shows that acquired prior information is not the only factor affecting the pattern of errors as a function of the true ball location. In addition to the bimodal prior effect, a linear relationship between errors and the true ball location was found. This effect can be well explained by biomechanical constraints and associated motor costs of potential movements (e.g. Griessbach et al.^32^). When the ball is close to the participant’s body (e.g. a ball position of 40 cm), participants generally tend to hit too far to the right, leading to a negative error, particularly when the opponent serves at a high speed. Data from day 1 (without the bimodal prior effect) and the control experiment suggest that balls played to the right end of the scale are most naturally hit by swinging the racket with an almost stretched arm—that is, a movement with low motor costs. When the ball is played closer to the body, participants have to flex their arm or make a small step to the left. The finding that movements are biased towards lower motor costs is well-established in the literature^31,33–36^. Consequently, in contrast to experiments that test Bayesian integration in isolated tasks, examining behaviour in highly complex tasks— such as returning tennis serves—highlights that movements are not only biased by prior expectations but also by other factors, such as biomechanical constraints. Importantly, however, the non-linear effects found in our data cannot be explained by biomechanical factors and, thus, show that humans do learn a bimodal prior and combine it with sensory information in action. Together, participants’ behaviour is well explained by a combined effect of the acquired bimodal prior and motor costs.

The results of the current study strengthen two decades of experimental lab-based work in perception^12^, cognition^13^, and motor control^5^ by demonstrating that Bayesian principles generalise to more complex cases including naturalistic movements and bimodal distributions. Specifically, regarding the learning of bimodal priors, the absence of any prior effect on day 1 is in line with previous studies that suggest that humans are, in principle, able to perform probabilistic inference with bimodal priors but require extensive experience to learn them^7,27^. More generally, the study puts into practice current calls to test major theories in complex, naturalistic behaviour^15–17^. In this regard, our work demonstrates that Bayesian theory provides a principled explanation of how our sensorimotor system solves impressive challenges at the limits of human performance, such as returning a tennis serve at over 250 km/h. This extension is particularly valuable for applied fields—from high-level sports to working environments or rehabilitation settings—suggesting that Bayesian theory provides a well-founded framework for addressing real-world challenges and, ultimately, advancing practice.

## Data availability statements

All raw data and prepared data used for statistical analysis and figure generation are available on GitHub (https://github.com/ispw-unibe-ch/bt-bimodal_prior_integration_vr_tennis).

## Code availability statements

The Python script for preparing the raw data, the Jupyter notebook for Bayesian simulations, and all R-scripts used for statistical analysis and figure generation are available on GitHub (https://github.com/ispw-unibe-ch/bt-bimodal_prior_integration_vr_tennis)

## Acknowledgements

The authors would like to thank Ralf Kredel, Martin Widmer, and Simon Maurer, as well as the Technology Platform of the Faculty of Human Sciences at the University of Bern, for the technical support.

## Author contributions

As co-first authors, S.Z. and D.B. contributed equally to the work. S.Z. came up with the experimental idea. S.Z., D.B., E.-J.H., and K.K. collaborated on further conceptualisation as well as on writing. D.B., S.Z. developed the experimental setup. D.B. led the data collection, and S.Z. and D.B. analysed the data.

## Methods

### Participants

24 healthy right-handed participants (13 females and 11 males; *M*_age_ = 22.0 years, *SD* = 2.2 years) with no distinctive tennis experience participated in the main experiment. 24 additional healthy right-handed participants (14 females and 10 males; *M*_age_ = 19.9 years, *SD* = 1.4 years) participated in the control experiment. The study protocol was approved by the ethics committee of the Faculty of Human Sciences at the University of Bern (approval number: 2017-12-00003) and was conducted in accordance with the Declaration of Helsinki. Written informed consent was obtained from all participants. The subject that appeared in the video of the experimental task provided written consent for publication.

### Virtual reality setup

Participants performed tennis returns in a custom life-sized VR CAVE environment (video on GitHub https://github.com/ispw-unibe-ch/bt-bimodal_prior_integration_vr_tennis). The 3D virtual tennis environment was developed using Unreal Engine 4.27 and is displayed stereoscopically in real-time and rendered in high resolution (pixel size 2.35 mm) using 11 projectors (Barco F50, 2560 ×1600 pixels, 60 Hz) driven by 11 cluster workstations on a 6.00 m × 3.75 m front wall, two 11.00 m × 3.75 m side walls, and a 6.00 m × 11.00 m floor. The head positions of the participants were tracked with an Optitrack 3D-Motion-Capture system at a rate of 200 Hz. The data were processed in real-time to ensure the accurate rendering of perspective from the participant’s point of view.

Participants only saw their real racket, while the virtual racket (of the same size and shape), which actually interacted with the virtual ball, was not displayed. The racket position was recorded using Optitrack (200 Hz). To provide haptic vibration feedback when the virtual ball was hit, a Wii controller was built into the handle of the racket (Figure 2, centre). To ensure synchronicity of the data streams, all the systems were coordinated using Streamix, an in-house development (https://tpf.philhum.unibe.ch/portfolio/streamix) based on the theoretical work by Maurer^37^.

The ball trajectories in Unreal Engine followed a prespecified spline trajectory (see Figure 2, left). To vary the serve location in the experiment, the spline of the serve was rotated around the vertical axes. In all speed conditions, the ball followed the same spline trajectory. We only manipulated how fast the ball followed the same spline trajectory, which does not reflect real physics but has the important advantage that the visual input only varied with regard to the time available to accumulate information. We also controlled that the temporal timing demands were the same in all speed conditions. Unbeknownst to the participants, they were actually striking an invisible stationary at a fixed location along the trajectory. This stationary ball could be hit within a 400-ms window—200 ms before and 200 ms after the visibly displayed ball passed its location.

This task was feasible but challenging. On average, participants hit 85.9% of the returns. The estimation error (horizontal deviation racket centre vs. ball) was also recorded when participants missed the ball. To achieve this, we implemented a second invisible plane, which was substantially larger than the real racket. When the virtual ball came into contact with this plane, an event was registered and the positions were recorded. However, the ball did not bounce off this plane, thus preserving the realism of the task. The only case in which no estimation error was recorded was when participants did not contact the ball within the 400-ms time window. Furthermore, when participants initiated their movement very late, it was possible that they hit the ball during a backswing movement. In these cases, participants were clearly too late to hit the ball over the net and the registered event did not represent their estimation error. Thus, these cases were identified as invalid and excluded from the data analysis (proportion of trials in slow condition: 0.40%, moderate condition: 1.24%, fast condition: 4.95%).

To provide a clearly defined task goal, a target was set on the opponent’s side of the tennis court (2.05 m in front of the service line and 2.05 m from the centre line; see Figure 1, left). Task performance (hitting the target) was directly linked to our main measure (horizontal deviation between the racket’s sweet spot and ball on the axes parallel to the tennis ground line at the moment of the impact: the ‘estimation error’). The estimation error scaled the direction of a prespecified spline trajectory. If the ball was perfectly hit with the sweet spot of the racket (zero deviation), the ball was directed to the centre of the target (see Figure 2, right). A deviation between the sweet spot and the ball at impact resulted in a deviation on the target by a factor of 20. In other words, a 1-cm deviation on the racket resulted in a 20-cm deviation in the minimum distance to the target. Thus, the estimation error determined how many points were awarded for the return. After each return, the points were displayed on a scale from 0 to 100. For a deviation of 15.5 cm and more (over the racket edge), participants received 0 points. For a perfect hit, they got 100 points; the points were linearly scaled in between. The impact angle and direction of the racket had no effect on the return trajectory. However, participants had to hit the ball at a minimum racket speed of 4 m/s. Below this threshold, the ball did not reach the opponent’s court over the net, which resulted in 0 points.

### Experimental design and procedure

In a within-subjects design, participants completed 3 experimental sessions within a week, with a minimum interval of 24 hours between sessions. Each session lasted approximately two hours. The participants’ task was to repeatedly return tennis serves. Their goal was to hit the centre of a target on the opponent’s side of the court (Figure 2, left) and, thereby, gain as many points as possible. To motivate the participants, a financial incentive was provided: the three participants with the highest scores over all three sessions received book vouchers (30, 50, and 80 Swiss francs).

At the beginning of each trial, participants’ initial position was standardised at 1.2 m behind the ground line and 3.0 m from the midline. Each trial began with the appearance of a red dot at the serving avatar’s position, accompanied by a brief acoustic signal. Following a random delay of 1 s–2 s, the virtual opponent initiated their serve. Importantly, while participants’ starting position before the trial was standardised, they were completely free to move as soon as they heard the acoustic signal.

All serves were played to the right of the participants’ starting position, and they returned the balls with a forehand strike. The distribution of the serve locations followed a bimodal distribution (mixture of Gaussians), with peaks at 50 cm (SD = 5 cm) and 90 cm (SD = 5 cm) to the right of the participants’ starting position (Figure 2, right).

We manipulated visual uncertainty by varying the initial ball speeds across three levels: slow (v = 108 km/h), moderate (v = 180 km/h), and fast (v = 252 km/h). While the trajectory of the approaching ball varied from trial to trial, the body movements of the serving avatar were identical for all serves. This implies that the opponent’s serving motion provided no cues regarding the ball’s trajectory. This design ensured that the participants could rely only on two sources of information: current sensory input regarding the ball’s trajectory and prior knowledge accumulated from previous trials.

Participants returned a total of 1,440 serves across 3 experimental sessions. Each session comprised 480 trials, organised into 10 blocks of 48 trials each. Within each block, the 48 trials included 16 trials per speed condition, with the speed conditions alternating in a predictable order (slow, moderate, and fast). Serve locations, however, were presented in a quasi-random order. The distribution of serve locations was controlled to remain identical across all days and speed conditions. Additionally, within each block and speed condition (16 trials), we ensured that the mean and standard deviation of serve locations were in a similar range. Specifically, for each side of the bimodal distribution (8 trials), the mean was no more than 1 cm away from one of the distribution peaks, and the standard deviation was no more than 2 cm higher than the overall standard deviation. On the first day, the participants had two additional warm-up blocks to familiarise themselves with the tennis task. The first warm-up block contained only slow serves and were all played to the same location (48 trials). In the second warm-up block, all serves were still played at a slow speed, but the locations were bimodally distributed as in a regular block (48 trials). Further details of the experimental protocols are available on GitHub (https://github.com/ispw-unibe-ch/bt-bimodal_prior_integration_vr_tennis). After the completion of the three experimental sessions, participants were asked whether they detected any pattern in the opponents’ serve distribution and if they recognised any technical facilitation (see Appendix).

### Predictions and analyses

#### Bayesian predictions

To generate predictions of participants’ responses in our task, we ran a Bayesian simulation according to Ma et al.^29^, which is documented in a step-by-step manner on GitHub (https://github.com/ispw-unibe-ch/bt-bimodal_prior_integration_vr_tennis). The model predicts participants’ responses when they estimate the ball position by combining likelihood *p(ball position_sensed_ |ball position_true_)* with prior knowledge *p(ball position_true_)* according to Bayesian integration. From Bayes’ rule, we obtain a posterior distribution:

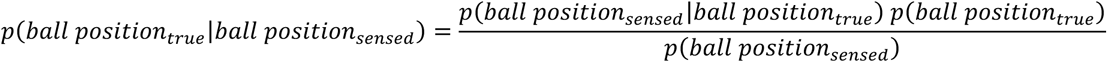

To estimate participants’ responses, we use the posterior mean estimate (PME) as a read-out, assuming that they try to optimise the mean squared error (MSE), as done in previous motor control studies^7^. For the bimodal distribution of true ball locations, the model predicts a non-linear relationship between the estimation error and the true ball position (see Figure 1). The simulations enabled the prediction of responses with different levels of uncertainty (the width of the likelihood function). Faster ball speeds are assumed to cause higher visual uncertainty (higher width of the likelihood function). The model predicts that the ‘jump’ between the left and right segments increases with higher uncertainty. However, as depicted in Figure 1, this happens only up to a certain point: When the uncertainty becomes extensively high, as could be the case with 252 km/h serves, the model prediction is that participants shift towards the mean of the distribution independently of the true ball position resulting in a linear relationship.

#### Biomechanical considerations

With the Bayesian model, we can predict systematic errors caused by integrating the learned prior distribution. However, in this task, it is reasonable to expect an additional systematic error as a function of true ball position caused by biomechanical constraints. Not every distance is reachable with equal effort (i.e. motor costs). We expected that swings would generally be biased towards distances with lower motor costs—in this case, distances where participants could swing their racket with an almost stretched arm. When the ball is played close to participants’ bodies, they need to flex their arms or take a step to the left, resulting in higher motor costs. The magnitude of the motor bias should depend on ball speed—while slow serves allow to compensate during the swing, it is much more difficult to do so for fast serves. This explanation results in a predictable pattern of errors: The error should linearly depend on the true ball position, with a steeper slope for higher ball speeds. This explanation was confirmed by data in the control experiment (Figure 3). However, this biomechanical explanation cannot explain non-linear relationships between the error and the true ball position. Non-linear relationships between the estimation error and the true ball position can only be effects of the bimodal prior.

#### Testing the bimodal prior effect: Multilevel regression analyses

To quantify the bimodal prior effect, we calculated regression models with two independent variables: true ball position and a bimodal segment factor (left vs. right). The dependent variable was the estimation error on the racket (see Figure 2, right).

The following procedures were used to check assumptions and handle missing values and outliers. First, we had multiple data points for each participant, and the residuals were not independent. Therefore, we built and compared regression models in several steps in order to take into account the hierarchical structure of the data^38^. Second, we checked the multicollinearity of the two independent factors (ball position and bimodal segment factor) with the variance inflation factor (VIF). Third, we graphically checked the homoscedasticity and normality of the residuals. Fourth, under the assumption that missing values occur randomly, multilevel regression analyses can handle missing values^38^. Fifth, we detected outliers using Cook’s distance and removed them when values affected the intercept more than three times as much as the mean.

In greater detail to the first point, the following steps of the construction of the models have been taken: We began with an intercept-only model, then added the ball position factor, the random intercepts, the random slopes, and, in the last step, the bimodal segment factor.

Descriptively, we compared models using the Akaike Information Criterion (AIC), Bayesian Information Criterion (BIC), and log-likelihood (Tables 2, 4, 6, 8, 10, and 12 in Extended Data). For inferential model comparisons, we used the log-likelihood ratio χ^2^ test. This enabled us to conclude whether the last added bimodal segment factor was a relevant and significant model extension in accordance with the parsimony principle^39^. Furthermore, we used the maximum likelihood for model estimations and tested regression coefficients with the Wald test for significance. For all inferential tests, we selected an alpha level of 5%. To account for multiple comparisons (three tests: slow, moderate, and fast), we used Bonferroni–Holm correction. For all statistics, we applied the R-package ‘nlme’^40^.

We analysed data from day 1 separately and from days 2 and 3 together, as previous research shows that learning a bimodal prior distribution needs to be consolidated over a night of sleep^30^. Thus, based on previous research, we expected no bimodal prior effect on day 1, while we expected an effect of the bimodal segment factor on days 2 and 3.

## Extended data

**Table 1.** Multilevel regression model of error estimation on day 1 in the fast condition.

| Fixed effects | *B* | *SE B* | 95% CI | *t*(2673) | *p* two-sided |
| --- | --- | --- | --- | --- | --- |
| Intercept | −18.32 | 4.01 | [−26.19, −10.46] | −4.57 | < .001 |
| Ball position | 0.16 | 0.06 | [0.03, 0.28] | 2.45 | .014 |
| Segment (0 = left, 1 = right) | 3.24 | 1.66 | [−0.01, 6.49] | 1.95 | .153 |
| Random effects |  |  |  |  |  |
| Intercept variance ($\tau_{00}$) | 280.98 | — | — | — | — |
| Slope variance ($\tau_{11}$) | 0.06 | — | — | — | — |
| Intercept-slope covariance ($\rho_{01}$) | −0.99 | — | — | — | — |
| Level-1 residual ($\sigma^2$) | 98.98 | — | — | — | — |
| ICC | 0.24 | — | — | — | — |

Note. *B* = unstandardized regression coefficients, *SE* = standard error, CI = confidence intervals, *t*(degrees of freedom). ICC = Interclass correlation coefficient. Model statistics: $N_{\text{participants}} = 24$ , $R^2_{\text{marginal}} = .148$ . Model comparison in Table 2.

**Table 2.** Model comparison for multilevel regression model of error estimation on day 1 in the fast condition.

| Model | *df* | AIC | BIC | logLik | Comparison | $\chi^2$ | *p* |
| --- | --- | --- | --- | --- | --- | --- | --- |
| 1 Intercept | 2 | 21171.00 | 21182.80 | −10583.50 |  |  |  |
| 2 Base model | 3 | 20755.30 | 20773.01 | −10374.65 | 1 vs 2 | 417.697 | < .001 |
| 3 Base model + RI | 4 | 20631.67 | 20655.28 | −10311.84 | 2 vs 3 | 125.632 | < .001 |
| 4 Base model + RS | 6 | 20211.17 | 20246.57 | −10099.58 | 3 vs 4 | 424.502 | < .001 |
| 5 Base model + RS +<br>Bimodal segment factor | 7 | 20209.35 | 20250.65 | −10097.67 | 4 vs 5 | 3.820 | .051 |

Note. *df* = degrees of freedom, AIC = Akaike Information Criterion, BIC = Bayesian Information Criterion, logLik = log-likelihood, RI = random intercepts, RS = random intercept and slopes. The base model includes the predictor ball position.

**Table 3.** Multilevel regression model of error estimation on day 1 in the moderate condition.

| Fixed effects | *B* | *SE B* | 95% CI | *t*(3096) | *p* two-sided |
| --- | --- | --- | --- | --- | --- |
| Intercept | −13.02 | 3.72 | [−20.30, −5.73] | −3.50 | < .001 |
| Ball position | 0.17 | 0.06 | [0.06, 0.28] | 3.04 | .002 |
| Segment (0 = left, 1 = right) | 0.27 | 1.46 | [−2.60, 3.15] | 0.19 | .852 |
| Random effects |  |  |  |  |  |
| Intercept variance ($\tau_{00}$) | 251.55 | — | — | — | — |
| Slope variance ($\tau_{11}$) | 0.04 | — | — | — | — |
| Intercept-slope covariance ($\rho_{01}$) | −0.96 | — | — | — | — |
| Level-1 residual ($\sigma^2$) | 90.90 | — | — | — | — |
| ICC | 0.29 | — | — | — | — |

Note. *B* = unstandardized regression coefficients, *SE* = standard error, CI = confidence intervals, *t*(degrees of freedom).
ICC = Interclass correlation coefficient. Model statistics: $N_{\text{participants}} = 24$ , $R^2_{\text{marginal}} = .090$ . Model comparison in Table 4.

**Table 4.** Model comparison for multilevel regression model of error estimation on day 1 in the moderate condition.

| Model | *df* | AIC | BIC | logLik | Comparison | $\chi^2$ | *p* |
| --- | --- | --- | --- | --- | --- | --- | --- |
| 1 Intercept | 2 | 24246.25 | 24258.34 | −12121.13 |  |  |  |
| 2 Base model | 3 | 23938.17 | 23956.31 | −11966.09 | 1 vs 2 | 310.078 | < .001 |
| 3 Base model + RI | 4 | 23570.14 | 22594.32 | −11781.07 | 2 vs 3 | 370.034 | < .001 |
| 4 Base model + RS | 6 | 23107.90 | 23144.18 | −11547.95 | 3 vs 4 | 466.239 | < .001 |
| 5 Base model + RS +<br>Bimodal segment factor | 7 | 23109.87 | 23152.19 | −11547.93 | 4 vs 5 | 0.035 | .852 |

Note. *df* = degrees of freedom, AIC = Akaike Information Criterion, BIC = Bayesian Information Criterion, logLik = log-likelihood, RI = random intercepts, RS = random intercept and slopes. The base model includes the predictor ball position.

**Table 5.** Multilevel regression model of error estimation on day 1 in the slow condition.

| Fixed effects | *B* | *SE B* | 95% CI | *t*(3049) | *p* two-sided |
| --- | --- | --- | --- | --- | --- |
| Intercept | 1.53 | 3.25 | [−4.84, 7.90] | 0.47 | .638 |
| Ball position | 0.08 | 0.05 | [−0.02, 0.18] | 1.62 | .105 |
| Segment (0 = left, 1 = right) | −2.24 | 1.39 | [−4.97, 0.49] | −1.61 | .216 |
| Random effects |  |  |  |  |  |
| Intercept variance ($\tau_{00}$) | 181.19 | – | – | – | – |
| Slope variance ($\tau_{11}$) | 0.03 | – | – | – | – |
| Intercept-slope covariance ($\rho_{01}$) | −0.94 | – | – | – | – |
| Level-1 residual ($\sigma^2$) | 79.84 | – | – | – | – |
| ICC | 0.29 | – | – | – | – |

Note. *B* = unstandardized regression coefficients, *SE* = standard error, CI = confidence intervals, *t*(degrees of freedom).
ICC = Interclass correlation coefficient. Model statistics: $N_{\text{participants}} = 24$ , $R^2_{\text{marginal}} = .003$ . Model comparison in Table 6.

**Table 6.** Model comparison for multilevel regression model of error estimation on day 1 in the slow condition.

| Model | *df* | AIC | BIC | logLik | Comparison | $\chi^2$ | *p* |
| --- | --- | --- | --- | --- | --- | --- | --- |
| 1 Intercept | 2 | 23242.37 | 23254.44 | −11619.19 |  |  |  |
| 2 Base model | 3 | 23229.23 | 23247.33 | −11611.62 | 1 vs 2 | 15.141 | < .001 |
| 3 Base model + RI | 4 | 22743.84 | 22767.97 | −11367.92 | 2 vs 3 | 487.399 | < .001 |
| 4 Base model + RS | 6 | 22366.15 | 22402.34 | −11177.08 | 3 vs 4 | 381.693 | < .001 |
| 5 Base model + RS + Bimodal segment factor | 7 | 22365.56 | 22407.78 | −11175.78 | 4 vs 5 | 2.595 | .108 |

Note. *df* = degrees of freedom, AIC = Akaike Information Criterion, BIC = Bayesian Information Criterion, logLik = log-likelihood, RI = random intercepts, RS = random intercept and slopes. The base model includes the predictor ball position.

**Table 7.**
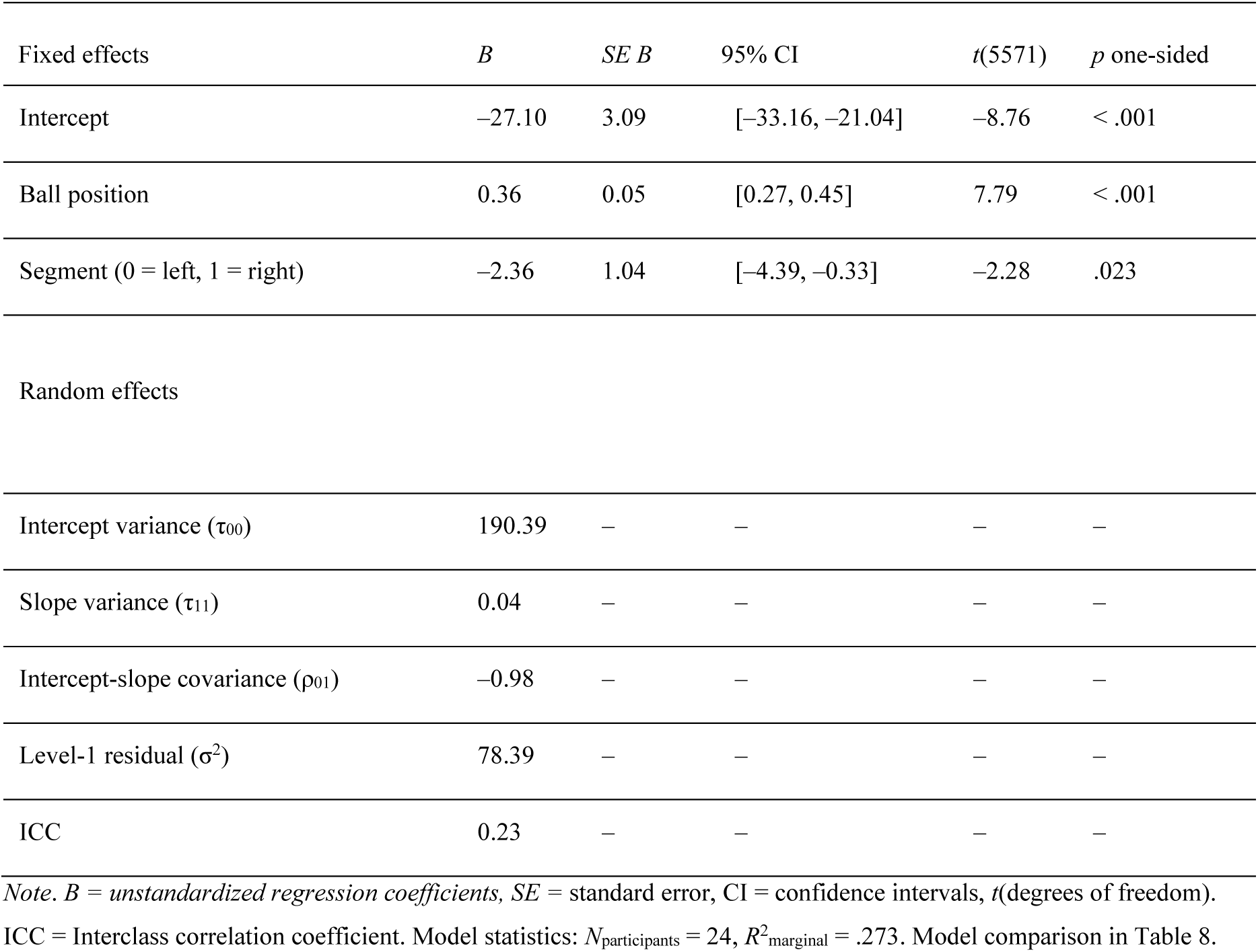
Multilevel regression model of error estimation on day 2+3 in the fast condition.

| Fixed effects | *B* | *SE B* | 95% CI | *t*(5571) | *p* one-sided |
| --- | --- | --- | --- | --- | --- |
| Intercept | −27.10 | 3.09 | [−33.16, −21.04] | −8.76 | < .001 |
| Ball position | 0.36 | 0.05 | [0.27, 0.45] | 7.79 | < .001 |
| Segment (0 = left, 1 = right) | −2.36 | 1.04 | [−4.39, −0.33] | −2.28 | .023 |
| Random effects |  |  |  |  |  |
| Intercept variance ($\tau_{00}$) | 190.39 | — | — | — | — |
| Slope variance ($\tau_{11}$) | 0.04 | — | — | — | — |
| Intercept-slope covariance ($\rho_{01}$) | −0.98 | — | — | — | — |
| Level-1 residual ($\sigma^2$) | 78.39 | — | — | — | — |
| ICC | 0.23 | — | — | — | — |

Note. *B* = unstandardized regression coefficients, *SE* = standard error, CI = confidence intervals, *t*(degrees of freedom).
ICC = Interclass correlation coefficient. Model statistics: $N_{\text{participants}} = 24$ , $R^2_{\text{marginal}} = .273$ . Model comparison in Table 8.

**Table 8.** Model comparison for multilevel regression model of error estimation on day 2+3 in the fast condition.

| Model | *df* | AIC | BIC | logLik | Comparison | $\chi^2$ | *p* |
| --- | --- | --- | --- | --- | --- | --- | --- |
| 1 Intercept | 2 | 43268.33 | 43281.59 | −21632.17 |  |  |  |
| 2 Base model | 3 | 41641.60 | 41661.49 | −20817.80 | 1 vs 2 | 1628.732 | < .001 |
| 3 Base model + RI | 4 | 41273.01 | 41273.53 | −20619.51 | 2 vs 3 | 396.587 | < .001 |
| 4 Base model + RS | 6 | 40521.26 | 40521.04 | −20234.63 | 3 vs 4 | 769.756 | < .001 |
| 5 Base model + RS + Bimodal segment factor | 7 | 40524.07 | 40524.48 | −20232.04 | 4 vs 5 | 5.184 | .023 |

Note. *df* = degrees of freedom, AIC = Akaike Information Criterion, BIC = Bayesian Information Criterion, logLik = log-likelihood, RI = random intercepts, RS = random intercept and slopes. The base model includes the predictor ball position.

**Table 9.** Multilevel regression model of error estimation on day 2+3 in the moderate condition.

| Fixed effects | *B* | *SE B* | 95% CI | *t*(6584) | *p* one-sided |
| --- | --- | --- | --- | --- | --- |
| Intercept | −18.20 | 2.90 | [−24.08, −12.70] | −6.28 | < .001 |
| Ball position | 0.29 | 0.04 | [0.21, 0.37] | 7.26 | < .001 |
| Segment (0 = left, 1 = right) | −2.64 | 0.91 | [−4.36, −0.94] | −2.90 | .006 |
| Random effects |  |  |  |  |  |
| Intercept variance ($\tau_{00}$) | 171.07 | – | – | – | – |
| Slope variance ($\tau_{11}$) | 0.03 | – | – | – | – |
| Intercept-slope covariance ($\rho_{01}$) | −0.97 | – | – | – | – |
| Level-1 residual ($\sigma^2$) | 71.51 | – | – | – | – |
| ICC | 0.24 | – | – | – | – |

Note. *B* = unstandardized regression coefficients, *SE* = standard error, CI = confidence intervals, *t*(degrees of freedom). ICC = Interclass correlation coefficient. Model statistics: $N_{\text{participants}} = 24$ , $R^2_{\text{marginal}} = .182$ . Model comparison in Table 10.

**Table 10.** Model comparison for multilevel regression model of error estimation on day 2+3 in the moderate condition.

| Model | *df* | AIC | BIC | logLik | Comparison | $\chi^2$ | *p* |
| --- | --- | --- | --- | --- | --- | --- | --- |
| 1 Intercept | 2 | 50105.85 | 50119.45 | −25050.93 |  |  |  |
| 2 Base model | 3 | 48784.41 | 48804.80 | −24389.21 | 1 vs 2 | 1323.442 | < .001 |
| 3 Base model + RI | 4 | 48022.77 | 48049.96 | −24007.39 | 2 vs 3 | 763.638 | < .001 |
| 4 Base model + RS | 6 | 47179.99 | 47220.77 | −23584.00 | 3 vs 4 | 846.782 | < .001 |
| 5 Base model + RS +<br>Bimodal segment factor | 7 | 47173.56 | 47221.13 | −23579.78 | 4 vs 5 | 8.435 | .004 |

Note. *df* = degrees of freedom, AIC = Akaike Information Criterion, BIC = Bayesian Information Criterion, logLik = log-likelihood, RI = random intercepts, RS = random intercept and slopes. The base model includes the predictor ball position.

**Table 11.** Multilevel regression model of error estimation on day 2+3 in the slow condition.

| Fixed effects | *B* | *SE B* | 95% CI | *t*(6462) | *p* one-sided |
| --- | --- | --- | --- | --- | --- |
| Intercept | −0.95 | 2.34 | [−5.53, 3.64] | −0.40 | .686 |
| Ball position | 0.10 | 0.03 | [0.03, 0.16] | 2.79 | .005 |
| Segment (0 = left, 1 = right) | −0.81 | 0.80 | [−2.38, 0.76] | −1.02 | .154 |
| Random effects |  |  |  |  |  |
| Intercept variance ($\tau_{00}$) | 107.93 | – | – | – | – |
| Slope variance ($\tau_{11}$) | 0.02 | – | – | – | – |
| Intercept-slope covariance ($\rho_{01}$) | −0.96 | – | – | – | – |
| Level-1 residual ($\sigma^2$) | 54.89 | – | – | – | – |
| ICC | 0.24 | – | – | – | – |

Note. *B* = unstandardized regression coefficients, *SE* = standard error, CI = confidence intervals, *t*(degrees of freedom). ICC = Interclass correlation coefficient. Model statistics: $N_{\text{participants}} = 24$ , $R^2_{\text{marginal}} = .033$ . Model comparison in Table 12.

**Table 12.** Model comparison for multilevel regression model of error estimation on day 2+3 in the slow condition.

| Model | *df* | AIC | BIC | logLik | Comparison | $\chi^2$ | *p* |
| --- | --- | --- | --- | --- | --- | --- | --- |
| 1 Intercept | 2 | 46272.21 | 46285.77 | −23134.11 |  |  |  |
| 2 Base model | 3 | 46045.41 | 46065.74 | −23019.70 | 1 vs 2 | 228.806 | < .001 |
| 3 Base model + RI | 4 | 45356.42 | 45383.53 | −22674.21 | 2 vs 3 | 690.987 | < .001 |
| 4 Base model + RS | 6 | 44591.78 | 44632.44 | −22289.89 | 3 vs 4 | 768.645 | < .001 |
| 5 Base model + RS + Bimodal segment factor | 7 | 44592.74 | 44640.19 | −22289.37 | 4 vs 5 | 1.034 | .309 |

Note. *df* = degrees of freedom, AIC = Akaike Information Criterion, BIC = Bayesian Information Criterion, logLik = log-likelihood, RI = random intercepts, RS = random intercept and slopes. The base model includes the predictor ball position.

**Table 13.** Multilevel regression model of error estimation for the control experiment in the fast condition.

| Fixed effects | *B* | *SE B* | 95% CI | *t*(2738) | *p* two-sided |
| --- | --- | --- | --- | --- | --- |
| Intercept | −31.69 | 2.72 | [−37.02, −26.35] | −11.65 | < .001 |
| Ball position | 0.38 | 0.04 | [0.31, 0.45] | 10.80 | < .001 |
| Random effects |  |  |  |  |  |
| Intercept variance ($\tau_{00}$) | 156.34 | — | — | — | — |
| Slope variance ($\tau_{11}$) | 0.03 | — | — | — | — |
| Intercept-slope covariance ($\rho_{01}$) | −0.91 | — | — | — | — |
| Level-1 residual ($\sigma^2$) | 129.81 | — | — | — | — |
| ICC | 0.21 | — | — | — | — |

Note. *B* = unstandardized regression coefficients, *SE* = standard error, CI = confidence intervals, *t*(degrees of freedom).
ICC = Interclass correlation coefficient. Model statistics: $N_{\text{participants}} = 24$ , $R^2_{\text{marginal}} = .207$ . Model comparison in Table 14.

**Table 14.** Model comparison for multilevel regression model of error estimation for the control experiment in the fast condition.

| Model | *df* | AIC | BIC | logLik | Comparison | $\chi^2$ | *p* |
| --- | --- | --- | --- | --- | --- | --- | --- |
| 1 Intercept | 2 | 22636.72 | 22648.57 | −11316.36 |  |  |  |
| 2 Base model | 4 | 21959.43 | 21977.20 | −10976.71 | 1 vs 2 | 679.295 | < .001 |
| 3 Base model + RI | 5 | 21539.11 | 21562.80 | −10765.55 | 2 vs 3 | 422.321 | < .001 |
| 4 Base model + RS | 7 | 21423.08 | 21458.62 | −10705.54 | 3 vs 4 | 120.028 | < .001 |

Note. *df* = degrees of freedom, AIC = Akaike Information Criterion, BIC = Bayesian Information Criterion, logLik = log-likelihood, RI = random intercepts, RS = random intercept and slopes. The base model includes the predictor ball position.

**Table 15.** Multilevel regression model of error estimation for the control experiment in the moderate condition.

| Fixed effects | *B* | *SE B* | 95% CI | *t*(3018) | *p* two-sided |
| --- | --- | --- | --- | --- | --- |
| Intercept | −13.13 | 2.40 | [−17.83, −8.42] | −5.46 | < .001 |
| Ball position | 0.15 | 0.03 | [0.09, 0.21] | 4.93 | < .001 |
| Random effects |  |  |  |  |  |
| Intercept variance ($\tau_{00}$) | 124.11 | — | — | — | — |
| Slope variance ($\tau_{11}$) | 0.02 | — | — | — | — |
| Intercept-slope covariance ($\rho_{01}$) | −0.88 | — | — | — | — |
| Level-1 residual ($\sigma^2$) | 100.02 | — | — | — | — |
| ICC | 0.25 | — | — | — | — |

Note. *B* = unstandardized regression coefficients, *SE* = standard error, CI = confidence intervals, *t*(degrees of freedom).
ICC = Interclass correlation coefficient. Model statistics: $N_{\text{participants}} = 24$ , $R^2_{\text{marginal}} = .050$ . Model comparison in Table 16.

**Table 16.** Model comparison for multilevel regression model of error estimation for the control experiment in the moderate condition.

| Model | *df* | AIC | BIC | logLik | Comparison | $\chi^2$ | *p* |
| --- | --- | --- | --- | --- | --- | --- | --- |
| 1 Intercept | 2 | 23660.52 | 23672.56 | −11828.26 |  |  |  |
| 2 Base model | 4 | 23465.21 | 23483.28 | −11729.61 | 1 vs 2 | 197.304 | < .001 |
| 3 Base model + RI | 5 | 22921.17 | 22945.26 | −11456.59 | 2 vs 3 | 546.042 | < .001 |
| 4 Base model + RS | 7 | 22798.38 | 22834.51 | −11393.19 | 3 vs 4 | 126.790 | < .001 |

Note. *df* = degrees of freedom, AIC = Akaike Information Criterion, BIC = Bayesian Information Criterion, logLik = log-likelihood, RI = random intercepts, RS = random intercept and slopes. The base model includes the predictor ball position.

**Table 17.** Multilevel regression model of error estimation for the control experiment in the slow condition.

| Fixed effects | *B* | *SE B* | 95% CI | *t*(2894) | *p* two-sided |
| --- | --- | --- | --- | --- | --- |
| Intercept | 7.22 | 2.18 | [2.96, 11.49] | 3.32 | .001 |
| Ball position | −0.05 | 0.02 | [−0.09, 0.00] | −1.96 | .050 |
| Random effects |  |  |  |  |  |
| Intercept variance ($\tau_{00}$) | 101.92 | – | – | – | – |
| Slope variance ($\tau_{11}$) | 0.01 | – | – | – | – |
| Intercept-slope covariance ($\rho_{01}$) | −0.88 | – | – | – | – |
| Level-1 residual ($\sigma^2$) | 81.31 | – | – | – | – |
| ICC | 0.26 | – | – | – | – |

Note. *B* = unstandardized regression coefficients, *SE* = standard error, CI = confidence intervals, *t*(degrees of freedom).
ICC = Interclass correlation coefficient. Model statistics: $N_{\text{participants}} = 24$ , $R^2_{\text{marginal}} = .006$ . Model comparison in Table 18.

**Table 18.** Model comparison for multilevel regression model of error estimation for the control experiment in the slow condition.

| Model | *df* | AIC | BIC | logLik | Comparison | $\chi^2$ | *p* |
| --- | --- | --- | --- | --- | --- | --- | --- |
| 1 Intercept | 2 | 21976.13 | 21988.09 | −10986.07 |  |  |  |
| 2 Base model | 3 | 21963.32 | 21981.26 | −10978.66 | 1 vs 2 | 14.810 | < .001 |
| 3 Base model + RI | 4 | 21364.30 | 21370.22 | −10669.15 | 2 vs 3 | 619.024 | < .001 |
| 4 Base model + RS | 6 | 21263.17 | 21299.04 | −10625.58 | 3 vs 4 | 87.130 | < .001 |

Note. *df* = degrees of freedom, AIC = Akaike Information Criterion, BIC = Bayesian Information Criterion, logLik = log-likelihood, RI = random intercepts, RS = random intercept and slopes. The base model includes the predictor ball position.

## Appendix

At the end of the last experimental session, participants were asked a few questions regarding the experiment. The questionnaire contained the following questions (translated from German):

1. Did you notice a pattern regarding the locations of the opponent’s serves?

2. Draw where you hit the balls with a cloud of dots.

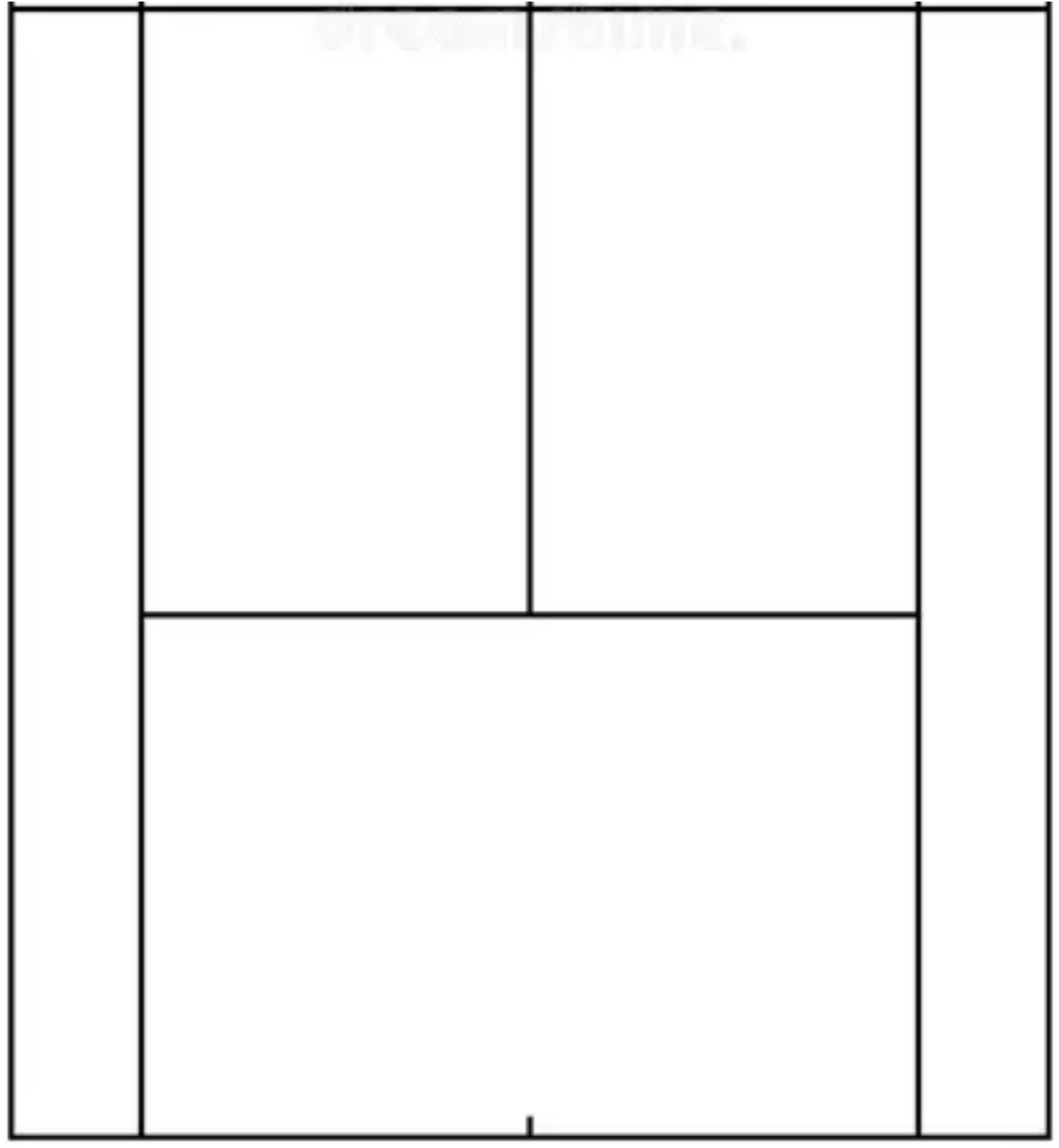

3. Circle the most appropriate answer:

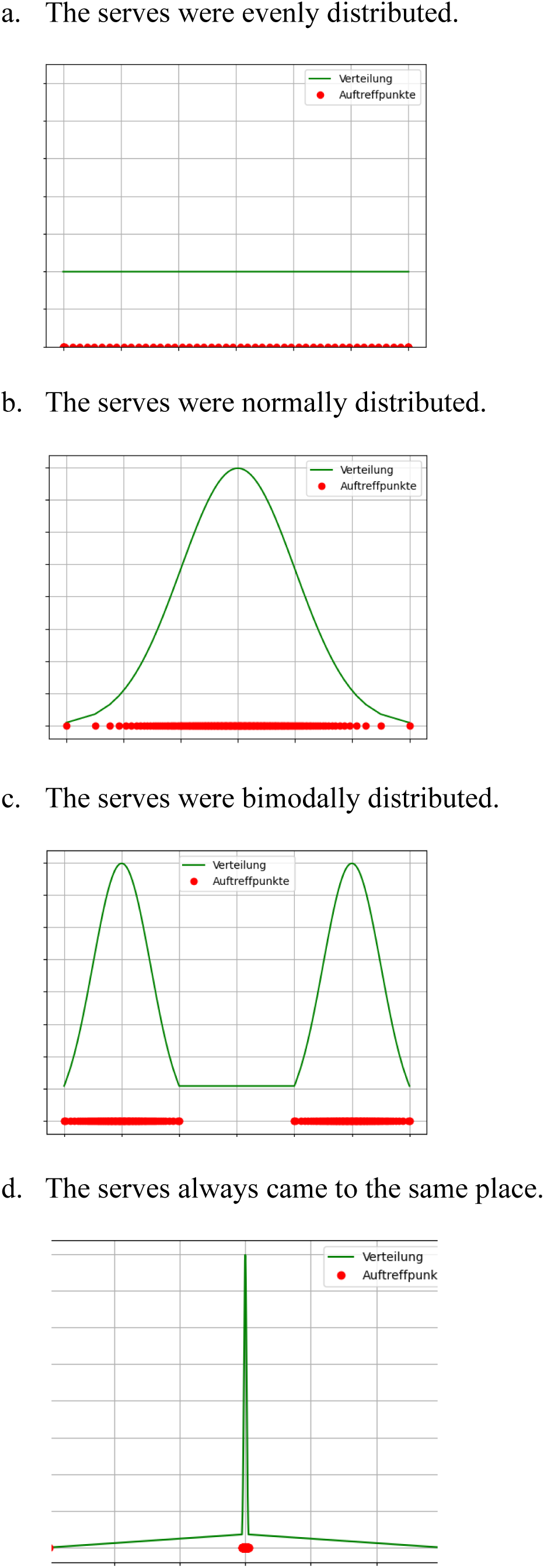

4. How sure are you about your answer?

Indicate as a percentage how sure you are of your answer:

5. Returning the serves was technically facilitated. Did you notice anything? Did you notice how? Please write down everything you noticed:

